# Effects of diffusion signal modeling and segmentation approaches on subthalamic nucleus parcellation

**DOI:** 10.1101/2021.02.28.433251

**Authors:** Demetrio Milardi, Gianpaolo Antonio Basile, Joshua Faskowitz, Salvatore Bertino, Angelo Quartarone, Giuseppe Anastasi, Alessia Bramanti, Alberto Cacciola

## Abstract

The subthalamic nucleus (STN) is commonly used as a surgical target for deep brain stimulation in movement disorders such as Parkinson’s Disease. Tractography-derived connectivity-based parcellation (CBP) has been recently proposed as a suitable tool for non-invasive in vivo identification and pre-operative targeting of specific functional territories within the human STN. However, a well-established, accurate and reproducible protocol for STN parcellation is still lacking. The present work aims at testing the effects of different tractography-based approaches for the reconstruction of STN functional territories.

We reconstructed functional territories of the STN on the high-quality dataset of 100 unrelated healthy subjects and on the test-retest dataset of the Human Connectome Project (HCP) repository. Connectivity-based parcellation was performed with a hypothesis-driven approach according to cortico-subthalamic connectivity, after dividing cortical areas into three groups: associative, limbic and sensorimotor. Four parcellation pipelines were compared, combining different signal modeling techniques (single-fiber vs multi-fiber) and different parcellation approaches (winner takes all parcellation vs fiber density thresholding). We tested these procedures on STN regions of interest obtained from three different, commonly employed, subcortical atlases. We evaluated the pipelines both in terms of between-subject similarity, assessed on the cohort of 100 unrelated healthy subjects, and of within-subject similarity, using a second cohort of 44 subjects with available test-retest data. We found that each parcellation provides converging results in terms of location of the identified parcels, but with significative variations in size and shape. Higher between-subject similarity was found with multi-fiber signal modeling techniques combined with fiber density thresholding. All the pipelines obtained very high within-subject similarity, with tensor-based approaches outperforming multi-fiber pipelines. We suggest that a fine-tuning of tractography-based parcellation may lead to higher reproducibility and aid the development of an optimized surgical targeting protocol.

## 1. Introduction

The subthalamic nucleus (STN) is crucially involved in the basal ganglia circuitry. Mainly composed of glutamatergic neurons (Kitai and Kita, 1987; Smith and Parent, 1988) with few GABA-ergic interneurons (Lévesque and André, 2005), the STN sends glutamatergic efferences to the striatum, the internal portion of the globus pallidus (GPi) and the two subdivisions of substantia nigra, pars compacta (SNc) and pars reticulata (SNr), while its main afference is represented by the external portion of the globus pallidus (GPe) (Hazrati and Parent, 1992; Parent and Hazrati, 1995; Parent and Parent, 2007; Smith et al., 1994, 1990). In addition, direct glutamatergic projections from the cerebral cortex have also been described (Nambu et al., 2002, 2000, 1997, 1996).

As for many other structures of the basal ganglia network, connections to and from STN are topographically organized. This feature allows the identification of distinct, yet integrated functional territories within all the basal ganglia structures (Alexander et al., 1990; Milardi et al., 2019). STN can be thus subdivided according to its connectivity into three functional territories: a rostromedial limbic territory, a ventrolateral associative territory and a dorsolateral motor territory (Joel and Weiner, 1997; Karachi et al., 2005; Parent and Hazrati, 1995; Shink et al., 1996). A similar topographical organization is also evident for direct cortical projections: connections arising from the primary motor cortex and supplementary motor area target the dorsal STN (Nambu et al., 1997, 1996), while projections originating from the dorsolateral prefrontal cortex and anterior cingulate cortex terminate in the ventrolateral and medial STN respectively (Haynes and Haber, 2013).

With the increasing use of deep brain stimulation (DBS) in the treatment of Parkinson’s Disease (PD), interest in the functional anatomy of the STN has expanded. Suggested in late 90’s as an alternative to ablative treatments (Benazzouz et al., 1993; Krack et al., 1998, 1997; Limousin et al., 1995), STN-DBS has a proved efficacy in treating both motor symptoms of PD and levodopa-induced dyskinesias (Deuschl et al., 2006; Rodriguez-Oroz et al., 2005; Weaver et al., 2009). More recently, DBS of STN has also been suggested for the treatment of psychiatric conditions such as treatment-resistant obsessive-compulsive disorder (OCD) (Mallet et al., 2002; Mallet et al., 2008; Haynes and Mallet, 2010; Welter et al., 2011). Prior knowledge about the topographical organization of STN provides a strong background for understanding differential STN DBS effects; while positive effects of DBS in PD patients are thought to stem from targeting of the dorsal sensorimotor portion of STN (Akram et al., 2017), clinical benefits observed in OCD are likely to be related to the dorsolateral prefrontal cortex-ventrolateral STN connections (Tyagi et al., 2019).

Connectivity-based parcellation (CBP) based on diffusion tractography has been proposed as a useful tool for the identification of distinct functional territories within the STN (Hamani et al., 2017). CBP is an heterogeneous group of techniques that use connectivity data (e.g. from probabilistic tractography) to characterize the latent topographical organization of parcels within a given region of interest (ROI) (Eickhoff et al., 2015). In the last decade, CBP has been extensively used to identify spatially segregated territories within different basal ganglia structures (Bardinet et al., 2006; Cacciola et al., 2019c; Draganski et al., 2008; Tziortzi et al., 2014) and has been even used to explore effects of implant locations in DBS surgery in experimental pilot studies (da Silva et al., 2017; E. H. Middlebrooks et al., 2018; Patriat et al., 2018). Taken together these findings suggest that CBP may provide substantial contribution to clinicians for pre-surgical planning and targeting.

In the past years, several works applied CBP to the study of STN functional organization in the human brain *in vivo,* confirming the topographical arrangement of STN connectivity previously described in animals (Accolla et al., 2014; Avecillas-Chasin et al., 2015; Brunenberg et al., 2012; Lambert et al., 2012; Plantinga et al., 2018). Despite the body of work demonstrating the feasibility of reconstructing STN territories using probabilistic tractography, consistent methodological differences have yet to be quantified. The lack of a well-defined connectivity-based parcellation protocol, together with the very small sample sizes, can undermine the reproducibility of results and limit their possible translation into a clinical context.

Based on these simple premises, the aim of the present work is to compare the effects of different STN parcellation protocols in terms of between-subjects and within-subject reproducibility. We performed hypothesis-driven CBP of the human STN on two high-quality datasets (Van Essen et al., 2013): the first cohort of 100 unrelated healthy subjects was used to assess between-subjects variability, and the second cohort of 44 healthy subjects with available test-retest data to investigate within-subject reliability. We compared two different diffusion signal modeling: a multi-fiber approach based on multi-shell multi-tissue constrained spherical deconvolution (MSMT-CSD) (Jeurissen et al., 2014), and a single-fiber diffusion tensor-based method (DTI). In addition, two different parcellation strategies were compared: a winner-takes-all segmentation, and a fiber density thresholding approach. Therefore, four distinct parcellation pipelines, combining different signal modeling techniques (single-fiber vs multi-fiber) and different parcellation approaches (winner takes all parcellation vs fiber density thresholding), were compared on three different, commonly employed, STN atlases.

### 2. Materials and methods

#### 2.1. Subjects, data acquisition and preprocessing

High-quality structural and diffusion MRI data have been collected from the HCP repository. Data were acquired by the Washington University, University of Minnesota, and Oxford University (WU-Minn) HCP Consortium (Van Essen et al., 2013). The study was approved by the Washington University Institutional Review Board and informed consent was obtained from all subjects. All HCP subjects were scanned using a Siemens 3T Skyra scanner (Siemens Healthcare, Erlangen, Germany) customized with a Siemens SC72 gradient coil and stronger gradient power supply with maximum gradient amplitude of 100 mT/m with the aim of improving diffusion imaging. (Van Essen et al., 2012). Two datasets have been employed for the present work: the first dataset consisted of 100 unrelated healthy subjects (100UNR) (males = 46, females = 54; age range: 22–36 years), and the second dataset included 44 subjects with available test-retest MRI scans (TRT) (males = 13; females = 31; age range: 22-36 years). Notice that 5 subjects of the first group were also part of the TRT group. Structural scans included T1-weighted images, acquired with the following parameters: TE = 2.14 ms, TR = 2,400 ms, and voxel size = 0.7 mm (Ugurbil et al., 2013). A single-shot two-dimensional (2D) spin-echo multiband echo planar imaging (EPI) sequence was used to acquire diffusion weighted images (DWI). All DWIs were equally distributed over three shells (b values of 1,000, 2,000, and 3,000 s/mm^2^), with isotropic spatial resolution of 1.25 mm (Sotiropoulos et al., 2013). Data were available in a minimally preprocessed form, that includes correction of EPI susceptibility, eddy-current–induced distortions, gradient nonlinearities, subject motion, and within-subject co-registration of structural and diffusion images (Glasser et al., 2013).

#### 2.2. MRI post-processing

Both structural and diffusion images were post-processed in order to perform tractography. T1-weighted structural images underwent brain extraction and cortical and subcortical segmentation using BET, FAST and FIRST tools on FSL (Patenaude et al., 2011; Smith, 2002; Smith et al., 2004). The obtained masks were visually inspected and, if needed, modified by a trained neuroanatomist. T1-weighted images were also normalized to the 1mm version of MNI152 2006 brain template using affine and nonlinear registration (FNIRT toolbox) and direct and inverse transformations were saved. A 5-tissue segmented image, needed for the implementation of MSMT-CSD, was then obtained from the native-space datasets. The 5 tissue image, together with the DWI data, was used to run multi-shell multi-tissue CSD (MSMT-CSD), an improvement of CSD signal modelling technique, in which fiber Orientation Distribution Function (fODF) is estimated directly from deconvolution of DW signal with a reference single fiber response function (Tournier et al., 2008, 2007). The MSMT-CSD modelling technique represents a variant designed to support multi-shell data and to overcome classical CSD limitations when it comes to estimate fODF in presence of tissue type heterogeneity (Jeurissen et al., 2014). For the DTI analysis, a weighted linear least squares estimation of diffusion tensors was carried out (Veraart et al., 2013). All these steps, and the following tractography, were performed using the MrTrix3 software (http://www.mrtrix.org/) (Tournier et al., 2012).

#### 2.3. Tractography

A probabilistic whole-brain tractogram (WB) of 5 million streamlines was generated for each subject, both for the MSMT-CSD and for the tensor-based approach, using default tracking parameters.

1. For the MSMT-CSD approach, tractography was performed with the following options: algorithm = *iFOD2* (Tournier et al., 2010), step size = 0.625 mm (0.5 x voxel size), maximum angle = 45°, minimal fODF amplitude = 0.05.
2. For the DTI approach, tractography was performed with the following options: algorithm = *Tensor_Prob* (Jones, 2008), step size = 0.125 mm, maximum angle = 45°, fractional anisotropy (FA) cutoff value = 0.1.

#### 2.4. Regions of interest (ROIs) delineation

In the present work we opted for an atlas-guided identification of STN based on nonlinear registration of standard-space STN ROIs into each subject’s space. To account for the variability that may be introduced by the arbitrary choice of an atlas, we tested our pipelines on three different subcortical atlases: i) a high-field structural MRI-based atlas of the basal ganglia (ATAG) (Keuken et al., 2014); ii) a template-based structural atlas (CIT168-Reinforcement Learning) (Pauli et al., 2018) and a multi-modal atlas based on combination of histology, structural and diffusion MRI (DISTAL) (Ewert et al., 2018). Atlases were all made available in MNI ICBM 2009c standard space in the LeadDBS software (Horn et al., 2019). For each atlas, left and right STN ROIs were resliced to 1 mm voxel size and transformed to the MNI152 standard space using a freely available transformation (credited to Andreas Horn, https://dx.doi.org/10.6084/m9.figshare.3502238.v1), then to each subject’s native space using the inverse transformations obtained in the normalization step of structural preprocessing (see paragraph above).

Cortical ROI segmentation was performed on T1-weighted images using the FreeSurfer software (Fischl et al., 2002). Briefly, the process involves averaging of T1-weighted images, skull stripping, tessellation, topology correction and spherical inflation of the white matter surface (Fischl et al., 2002, 1999b, 1999a; Ségonne et al., 2004). A modified and improved version of FreeSurfer’s *recon-all* pipeline is part of the minimal preprocessing procedures provided by the HCP repository; further details can be found in Glasser et al. (2013). Parcellation of the cerebral cortex into regions with respect to gyral and sulcal structures was performed according to the Desikan-Killiany atlas (Desikan et al., 2006).

Like in previous studies (Cacciola et al., 2019c; Patriat et al., 2018; Plantinga et al., 2018), cortical gyral ROIs were merged into three function-related cortical targets: an associative target, consisting of superior, middle and inferior frontal gyri; a limbic target including lateral orbitofrontal cortex, medial orbitofrontal cortex, frontal pole and anterior cingulate cortex; and a sensorimotor target, corresponding to precentral gyrus, postcentral gyrus and paracentral lobule.

#### 2.5. Connectivity based parcellation

We performed CBP by applying the following pipeline, both to the MSMT-CSD and the DTI datasets:

1. From each 5-million-streamlines WB, tracts between cortical target ROIs and the STN were extracted. This step was carried out separately for each left and right STN ROI obtained with each of the three atlases, and versus each of the three cortical target ROIs (associative, limbic and sensorimotor) described above.
2. Each tractogram was converted into a track density image (TDI) (Calamante et al., 2010). In a TDI, intensity at each voxel is defined as the number of fiber tracts passing through a given grid element. Voxel size was fixed at 1 mm^3^, same as STN and all the other ROIs.
3. Then, each TDI was multiplied to the relative STN ROI, to retrieve connectivity density-weighted parcels on the STN volume.
4. To ensure that different connectivity would not affect the intensity values of each parcel, causing more connected parcels to have higher intensity values than less connected ones, each parcel was normalized by dividing each voxel’s intensity by the mean intensity of the whole parcel.

From this point on, the procedure was differentiated to obtain, for each dataset, two different kinds of parcellation: the first, by following a winner-takes-all (WTA) hard segmentation approach, i.e., by assigning each STN voxel to the parcel with highest connectivity density values compared with the others; the second, without using any explicit clustering method. For the WTA parcellation scheme, normalized parcels underwent hard segmentation by running the *find_the_biggest* command on FSL. For the parcellation scheme without WTA parcellation, a threshold was applied to the normalized parcels, to filter out voxels with lower connectivity density, that could confound the obtained results. In line with previous work, we used an arbitrary threshold of 25%, meaning that only voxels with track density higher than 25% of the whole track-density map were preserved (Plantinga et al., 2018). In summary, CBP resulted in four groups, representing diverging processing pipelines:

1. a CSD-tractography group with WTA segmentation as final step;
2. a CSD-tractography group with 25% fiber density thresholding;
3. a DTI-tractography group with WTA hard segmentation;
4. a DTI-tractography group with 25% fiber density thresholding;

from now on we will refer to these groups as CSD-WTA, CSD-thr25, DTI-WTA and DTI-thr25 respectively. Each of these pipelines was carried out separately on left and right STN ROIs obtained from each of the three atlases taken into account (DISTAL, ATAG and CIT168-RL), thus resulting in 24 possible combinations for each subject. Each pipeline subdivided the STN into three parcels (Associative, Limbic and Sensorimotor), driving to a total of 72 parcels for each subject.

Finally, all the obtained parcels were transformed to the MNI ICBM 2009b standard template.

For visualization purposes, the parcels obtained from the 100UNR group were binarized and summed up to obtain a maximum probability map (MPM) reflecting the average parcel at the whole sample level. A 50% threshold was applied to each MPM, i.e., only voxels that show overlap in at least half of the sample were taken into account. The entire processing pipeline is summarized in Figure 1.

**Figure 1.**
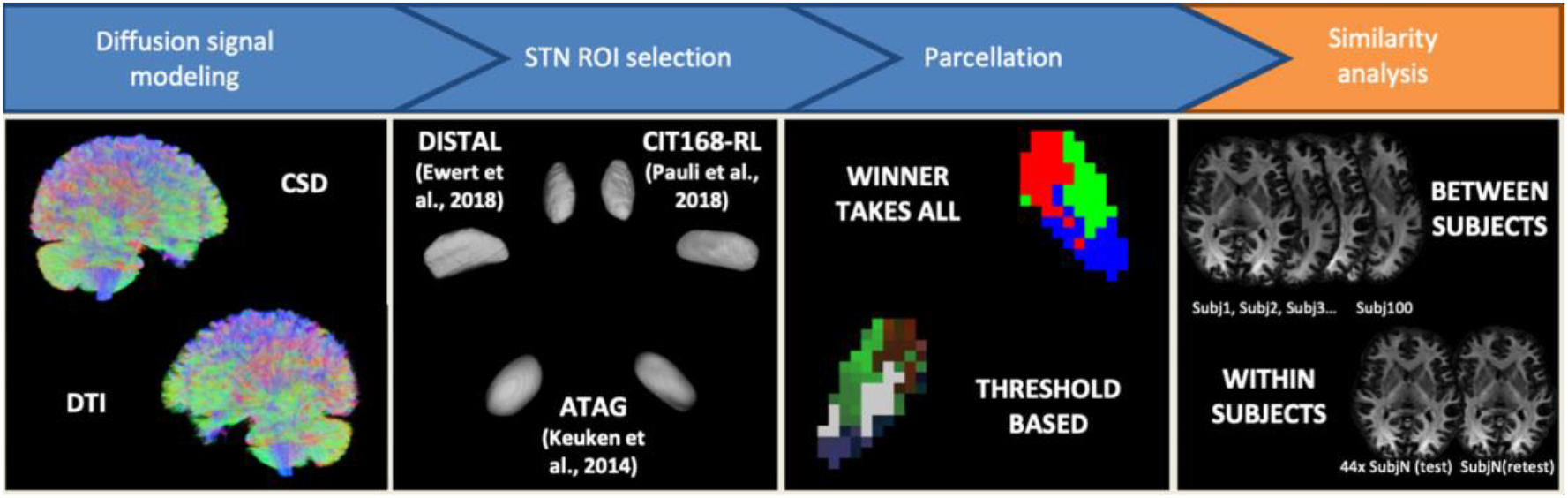
Methodological variables in STN parcellation. A summary of the processing pipeline, meant to highlight the methodological variables involved in our analysis. Separate results were generated for each combination of variables, and the outputs were tested in terms of both between- and within-subjects reliability. Full detail in text.

#### 2.6. Quantitative connectivity analysis

Quantitative connectivity for each target at subject level was estimated by calculating the streamline density index (SDI) (Cacciola et al., 2019c; Theisen et al., 2017), defined as the percentage ratio between each parcel volume and STN ROI volume:

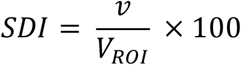

where *v* is the parcel volume in subject space and *V_ROI_* the STN ROI volume, in subject space. Statistically significant differences were assessed for each parcel grouped by side (left or right) and type (associative, limbic, sensorimotor). Specifically, a 3×4 repeated measures factorial ANOVA model was applied to investigate differences in SDI related to the use of different atlases (three levels: DISTAL, ATAG, CIT168-RL), the reconstruction pipeline employed (four levels: CSD-thr25, CSD-WTA, DTI-thr25, DTI-WTA) and their interaction. All the statistical analysis was carried out using SPSS Statistics (IBM SPSS Statistics for Windows, Version 25.0. Armonk, NY: IBM Corp.).

#### 2.7. Between-subject and within-subject similarity measures

Reproducibility of subthalamic parcellations was evaluated both in terms of between-subject similarity, that was assessed on the 100UNR cohort, and within-subject similarity, that was measured on the TRT data. Specifically, we evaluated reproducibility by calculating similarity indices for each parcel image after registration to the standard template; the result of these similarity indices is a number ranging between 0 and 1, where a value closer to 1 indicates higher similarity and a value closer to 0 indicates high dissimilarity.

Between-subject similarity was evaluated by calculating the overlap-by-label (OBL), a measure of the overlap of each parcel across all datasets, and the total accumulated overlap (TAO) that measures the overall, groupwise overlap for a given parcellation pipeline. Both TAO and OBL are based on the Tanimoto coefficient, which measures the similarity between different sets:

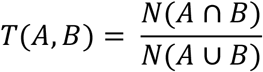

where N is expressed as number of voxels (Crum et al., 2006).

For a group of *m* pairs of images, where *m* represents all the possible pairwise combinations between images of the same parcel, OBL is defined as:

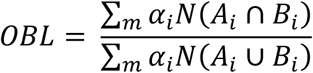

where *i* is the parcel label and α is a weighting coefficient. We defined α as the inverse of the mean of the absolute value of volumes for A and B, to avoid overestimation of larger parcels:

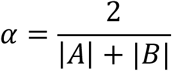

For the same group, TAO is defined as:

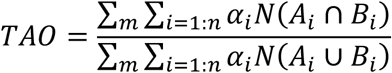

where *n* is the total number of parcels obtained from a given parcellation pipeline (CSD-thr25, CSD-WTA, DTI-thr25, DTI-WTA) (da Silva et al., 2017; Traynor et al., 2010). Both OBL and TAO were calculated separately for each hemisphere and for each of the three subcortical atlases taken into account.

Within-subject similarity was assessed by calculating the Dice similarity coefficient (DSC) (Crum et al., 2006), a popular similarity metric that is commonly used in neuroimaging studies. Dice similarity coefficient is defined as:

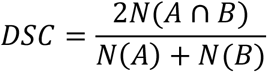

where A and B are images of the same parcel obtained for the same subject from the test and the retest scan, respectively, and N is expressed as number of voxels.

To investigate effects of the atlas choice and of the employed pipeline on within-subject similarity, we grouped individual DSC values for each hemisphere (left and right) and parcel type (associative, limbic and sensorimotor), and applied a 3×4 repeated measures factorial ANOVA using atlas and pipeline as within-subject factors.

#### 2.8 Spatial relations between center-of-gravity coordinates

For each subject, both in the 100UNR and in the TRT groups, the center-of-gravity for each subthalamic ROI was obtained using FSL’s *fslstats* command. This processed rendered an x, y, and z coordinate in standard space for each ROI of each pipeline. For the 100UNR group, we sought to characterize how these coordinates across 100 subjects varied based on parcellation strategy. We therefore obtained a 100-by-100 Euclidean distance matrix between center-of-gravity coordinates for each parcellation strategy (24 ROIs for each atlas: 72 strategies total). We then employed the Mantel test with a permutation approach (5000 permutations), to assess the statistical similarity of these distance matrices. The end result is a Mantel test conducted between each pair of pipelines, testing for the similarity of pairwise distance between strategies. For the TRT group, we sought to characterize the reliability of the center-of-gravity between test and retest scans. We therefore obtained a 44-by-44 test-retest Euclidean distance matrix between center-of-gravity coordinates of test and retest data, for each parcellation strategy (72 strategies total). Rows of this matrix documented subjects’ test data, whereas columns of this matrix represented subjects’ retest data. Therefore, these matrices were not symmetric. We rank transformed the rows of this matrix. For reliable data, we would expect the diagonal elements of this matrix to be near rank 1, indicating that subjects’ test coordinates are least distant from their respective re-test coordinates. We performed a permutation test (5000 permutations) by randomizing the ordering of the retest coordinates, to obtain the median diagonal rank expected by chance, and compared this to the empirical median diagonal rank to obtain a p-value.

### 3. Results

#### 3.1 Connectivity-based parcellation of the STN

For each subject, both in the 100UNR and in the TRT group, left and right STN were subdivided into three parcels using three different pairs of STN ROI and four different pipelines. Not all the pipelines were able to reconstruct all three parcels in the totality of subjects. In particular, CSD-based pipelines (CSD-thr25 and CSD-WTA) were able to reconstruct associative, limbic and sensorimotor STN parcels in both left and right hemisphere in 100 subjects (100%) of the 100UNR group, and 44 subjects (100%) of the TRT group both on test and retest scans, for each of the three distinct subcortical atlases. Conversely, the reconstruction of the right hemisphere associative parcel using DTI-based pipelines (DTI-thr25; DTI-WTA) failed in 1 subject (1%) of the 100UNR group, regardless of the employed subcortical atlas. The reconstruction of the right hemisphere sensorimotor parcel failed in 2 subjects (2%) of the 100UNR group for each of the three subcortical atlases, and in 1 subject (1%) only for the ATAG atlas. In the 100UNR group, limbic parcel was not successfully reconstructed in the left hemisphere in 17 subjects (17%) regardless of the employed subcortical atlas, in 2 subjects (2%) using ATAG and CIT168-RL atlases and in 21 subjects (21%) using ATAG atlas only; in the right hemisphere, reconstruction of the limbic parcel failed in 34 subjects regardless of the employed subcortical atlas, in 1 subject using DISTAL and ATAG atlases and in 11 subjects (11%) using ATAG atlas only. In the TRT group, the limbic parcel was not reconstructed for the left STN in 22 test datasets (50%) and 23 retest datasets (50.5%) regardless of the employed subcortical atlas, and in 6 test (13.6%) and 6 retest (13.6%) datasets using the ATAG atlas only; for the right STN, the reconstruction of the limbic parcel failed in 36 test (81.8%) and 37 retest (84%) datasets using each of the three employed subcortical atlases, and in 3 test (6.8%) and 4 retest (9.1%) using ATAG atlas only. Due to the high failure rate, the limbic parcel was excluded from all the statistical analyses. In all the described cases, failure was related to the tract selection step of the pipeline (see paragraph 2.5, step 1) as no streamlines connecting STN ROIs and target regions were extracted from the 5-million-streamlines WB. Related frequencies of successfully reconstructed parcels for each atlas and group are reported in Supplementary file 1.

For visualization purposes, the average population maps obtained from the 100UNR group are displayed in Figure 2. It is worth to note that, despite the evident differences in size and shape of STN ROIs across different subcortical atlases, STN parcels share a similar spatial organization: the associative parcel is located in the ventrolateral portion of STN, the limbic parcel in the ventromedial STN and sensorimotor parcel in the dorsolateral STN.

**Figure 2.**
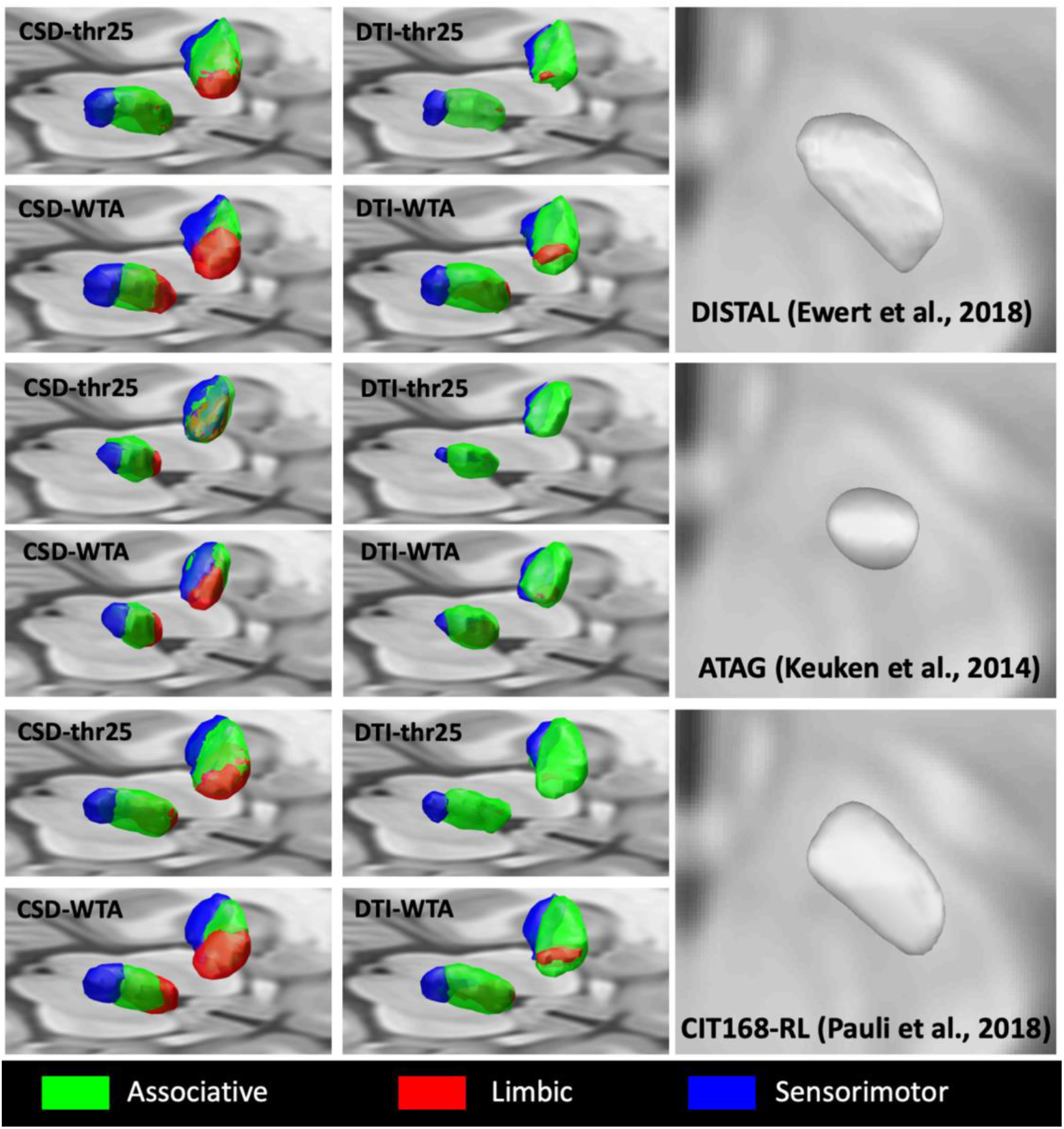
Group-level STN maps using different parcellation pipelines. 3D renderings of maximum probability maps retrieved after parcellation of the STN using different pipelines. Maximum probability maps are obtained after transformation of each subject map to template space; maps have been binarized and summed up across all subjects and a 50% population threshold was set. Hence, MPM volumes are representative of the number of voxels overlapping in at least 50% of the sample.

A similar spatial organization can be also observed by comparing the outputs of different parcellation pipelines within STN ROIs obtained from the same subcortical atlas (supplementary file 2). Thresholded MPMs of STN parcels show marked differences in volumes across different atlases and pipelines (Table 1).

**Table 1.** Volumes of maximum probability maps of STN (mm^3^). Maximum probability maps are obtained after transformation of each subject map to template space; maps have been binarized and summed up across all subjects and a 50% population threshold was set. Hence, MPM volumes are representative of the number of voxels overlapping in at least 50% of the sample.

| Atlas | Pipeline | L-Associative | L-Limbic | L-Sensorimotor | R-Associative | R-Limbic | R-Sensorimotor |
| --- | --- | --- | --- | --- | --- | --- | --- |
| **DISTAL**<br>(*Ewert et al., 2018*) | CSD-WTA | 222 | 201.375 | 209.5 | 227.625 | 215.25 | 206.875 |
|  | CSD-thr25 | 371 | 199.375 | 232.5 | 340.125 | 165.125 | 189.125 |
|  | DTI-WTA | 328.75 | 39.875 | 143 | 328 | 93.875 | 124.625 |
|  | DTI-thr25 | 200.75 | 10.375 | 88.625 | 209.5 | 2.375 | 67 |
| **ATAG**<br>(*Keuken et al., 2014*) | CSD-WTA | 166.75 | 145 | 172.5 | 167.5 | 143 | 163.5 |
|  | CSD-thr25 | 231 | 115.375 | 194.375 | 219.375 | 103.25 | 146.875 |
|  | DTI-WTA | 190.375 | 33.375 | 113.375 | 209.5 | 75.375 | 66 |
|  | DTI-thr25 | 122.75 | / | 61 | 130.125 | / | 30 |
| **CIT168-RL**<br>(*Pauli et al., 2018*) | CSD-WTA | 224.875 | 206.25 | 199.875 | 234.625 | 214.5 | 187 |
|  | CSD-thr25 | 364.875 | 193.875 | 208.5 | 341.5 | 167.125 | 168.5 |
|  | DTI-WTA | 339.5 | 49.875 | 146.375 | 350.5 | 98.875 | 120.25 |
|  | DTI-thr25 | 229 | 7.875 | 90 | 228.625 | 2.375 | 65 |

As reported in the “Materials and methods” section (paragraph 2.6), such volumes reflect the number of voxels overlapping in at least half of the sample after normalization to standard space. This number is generally higher for parcels obtained through CSD-based pipelines, in particular for the limbic parcels, that show very reduced or even absent voxel overlap in DTI-based pipelines, probably due to the high number of subjects in which such parcels were not reconstructed by using DTI.

#### 3.2 Quantitative connectivity analysis

To investigate how the choice of STN ROIs from different subcortical atlases and the use of different pipelines for parcellation influence the quantitative connectivity estimates, we conducted a 3×4 repeated measures ANOVA grouping parcels by hemisphere (left and right) and type (associative, sensorimotor).

For both left and right associative and sensorimotor parcels, Mauchly’s test of sphericity indicated a violation of sphericity for atlas, pipeline and their interaction factor. Degrees of freedom and p-values were adjusted using the Greenhouse-Geisser correction.

We found a significative effect of atlas choice on SDI values for both associative (left: F(1.10,108.96)=4388.37, p<0.001, η_p_^2^=0.98; right: F(1.10,108.42), p<0.001, η_p_^2^=0.97), and sensorimotor parcels (left: F(1.12,110.69)=1588.24, p<0.001, η_p_^2^=0.94; right: F(1.18,116.90)=1420.69, p<0.001, η_p_^2^=0.94).

A significative effect of pipeline selection was also found for all the analyzed parcels (left associative: F(2.30,227.17)=494.11, p<0.001, η_p_^2^=0.83; right associative: F(2.60,253.31)=399.23, p<0.001, η_p_^2^=0.80; left sensorimotor: F(1.96,194.00)=361.911, η_p_^2^=0.76; right sensorimotor: F(1.71,169.57)=528.22, p<0.001, η_p_^2^=0.84).

Finally, a significant interaction effect of atlas choice and pipeline selection was found for all parcels: left associative (F(2.68,265.25)=276.57, p<0.001, η_p_^2^=0.74), right associative (F(3.2,316.28)=284.34, p<0.001, η_p_^2^=0.74), left sensorimotor (F(2.33,230.93)=135.18, p<0.001, η_p_^2^=0.58) and right sensorimotor (F(2.56,253.87)=211.74, p<0.001, η_p_^2^=0.68). Distributions of SDI values across different parcels, grouped by atlas and pipeline, are plotted in Figure 3.

**Figure 3.**
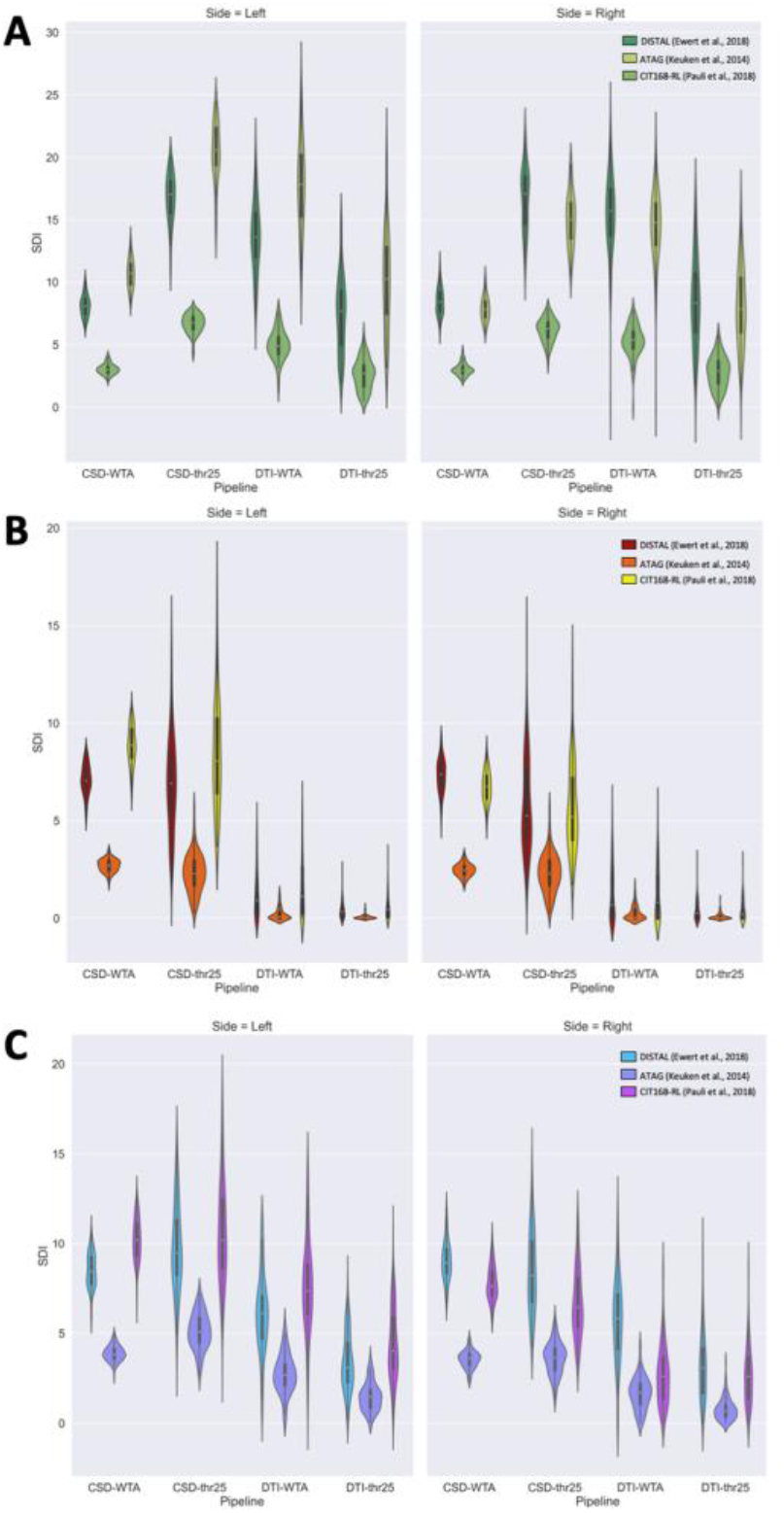
Subject-level volume differences across parcellation pipelines. Streamline Density Index (SDI) values for each parcellation pipeline are reported in violin plots. A) associative parcel; B) limbic parcel; C) sensorimotor parcel.

#### 3.3 Between-subjects and within-subject similarity

Between-subjects similarity was evaluated for each parcellation pipeline grouped by hemisphere and atlas using TAO, whose values are indicative of the groupwise accumulated overlap for all the parcels obtained using each pipeline. Independently of the atlas, we found a similar pattern such that higher TAO values were obtained using CSD-WTA (DISTAL atlas: left 0.65, right 0.66; ATAG atlas: left 0.67, right 0.67; CIT168-RL atlas: left 0.67; right 0.67) or CSD-thr25 (DISTAL atlas: left 0.67, right 0.65; ATAG atlas: left 0.68, right 0.66; CIT168-RL atlas: left 0.67, right 0.65), in comparison to the lower values obtained using DTI-WTA (DISTAL atlas: left 0.45, right 0.43; ATAG atlas: left 0.42; right 0.39; CIT168-RL atlas: left 0.46, right 0.43) or DTI-thr25 pipelines (DISTAL atlas: left 0.39, right 0.35; ATAG atlas: left 0.40, right 0.32; CIT168-RL atlas: left 0.40, right 0.36).

To investigate the relative weight of each parcel in driving the accumulated similarity values observed by TAO, we also evaluated similarity at the parcel-wise level by calculating OBL. The obtained values are plotted separately for each atlas in Figure 4A. Generally speaking, results confirm the trend observed in TAO values and related to CSD-based versus DTI-based parcellation, with the former performing better than the latter in most of the cases. Among different parcels, in most of the observed cases, the associative parcels scored the highest OBL values, followed by sensorimotor parcels, although few exceptions were observed, in particular using ATAG atlas. In all cases, limbic parcels obtained the lower OBL values. In addition, more marked differences were observed between WTA-based and threshold-based methods; in particular, for CSD-based pipelines, threshold-based approaches resulted in higher, but less uniform OBL values (range: 0.56-0.78), while WTA-based approaches obtained generally lower, but more uniform values (range: 0.63-0.71); conversely, for DTI-based pipelines, WTA-based pipelines always obtained higher OBL values (range: 0.10-0.73) in comparison to threshold-based pipelines (range: 0.10-0.55).

**Figure 4.**
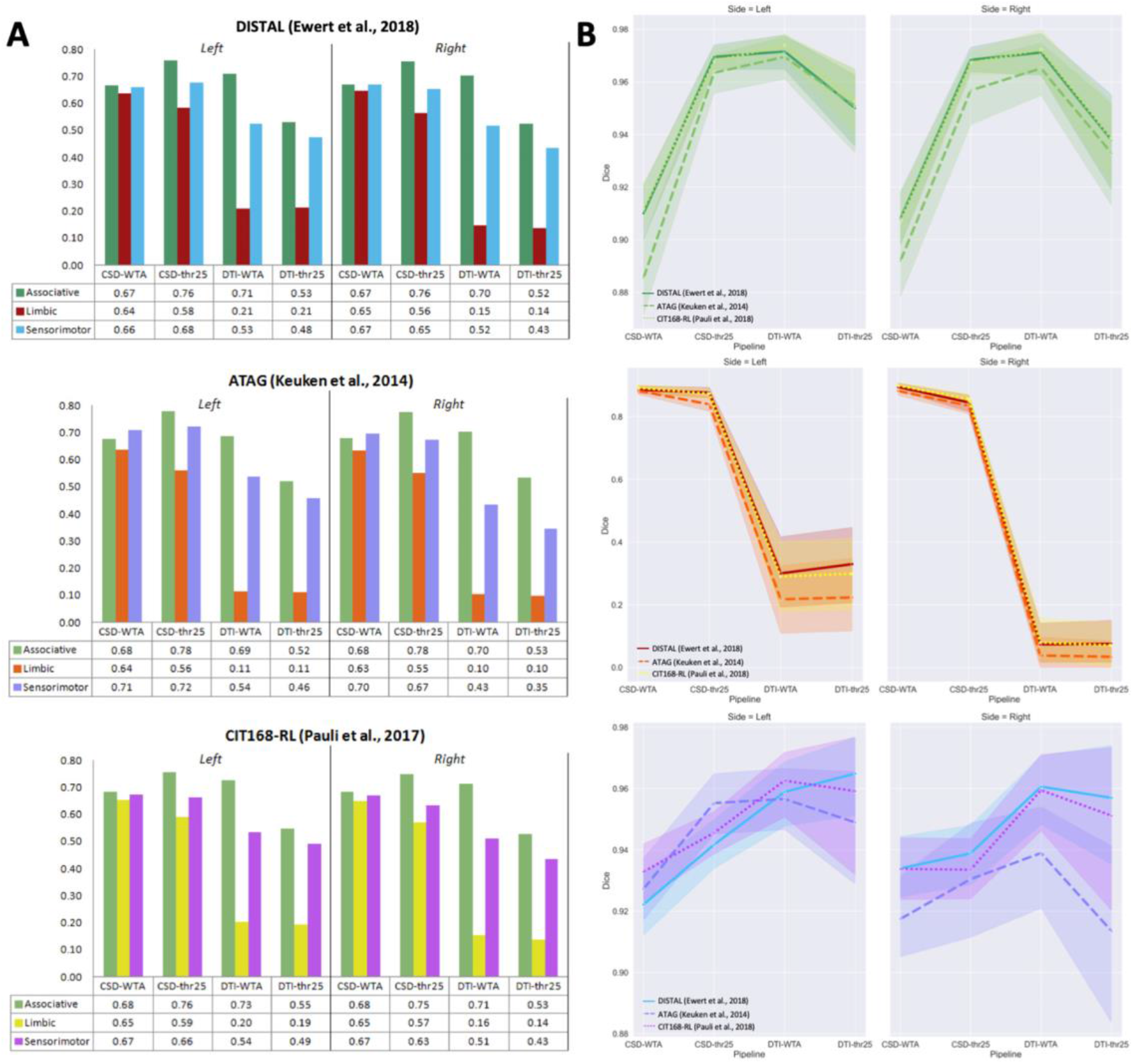
Between-subjects and within-subjects similarity measures. A) Overlap-by-label (OBL) values plotted for each parcel types. Values are reported in tables. B) Average Dice similarity coefficients (DSC) estimated on the test-retest (TRT) dataset, between parcels obtained with the same pipeline on test and retest data.

To evaluate and compare within-subject similarity across different atlases and pipelines, we calculated average DSC for each individual of the TRT datasets (Figure 4B). Then, we grouped DSC values by hemisphere (left and right) and parcel type (associative and sensorimotor) and conducted a 3×4 repeated measures ANOVA to investigate effects of atlas and pipeline choice.

Since Mauchly’s test of sphericity indicated a violation of sphericity for atlas, pipeline and their interaction factor for all parcels, degrees of freedom and p-values were adjusted using the Greenhouse-Geisser correction.

For left associative parcel, we found a significative effect of atlas (F(1.29,55.37)=9.742, p=0.001, η_p_^2^=0.19), pipeline (F(2.14,91.97)=79.44, p<0.001, η_p_^2^=0.65) and their interaction factor (F(3.82,164.37)=8.84, p<0.001, η_p_^2^=0.17). Conversely, for right associative parcel a significant effect on DSC was found both for atlas (F(1.25,53.83)=7.05, p=0.007, η_p_^2^=0.14) and pipeline choice (F(1.97,84.84)=64.92, p<0.001, η_p_^2^=0.60), but no significative interaction effects were found (p=0.35). A significative effect of atlas selection on DSC values was not found (p=0.56); conversely, a significative effect of pipeline choice (F(1.54,66.071)=64.92, p=0.001, η_p_^2^=0.18) and a significative interaction effect of atlas and pipeline choice (F(3.40,146.03)=64.92, p<0.019, η_p_^2^=0.070) were found. Finally, for right sensorimotor parcel, a significative effect was found for atlas (F(1.23,53.04)=14.57, p<0.001, η_p_^2^=0.25) and pipeline (F(1.98,89.18)=4.02, p=0.022, η_p_^2^=0.85) but no significative interaction effect was found (p=0.056).

#### 3.4 Variation between center-of-gravity coordinates between parcellation approaches

When assessing the variation between center-of-gravity coordinates between parcellation approaches, we observed significant similarities (corrected for false-discovery rate at α=0.01) between approaches (Figure 5A). These significant similarities indicate pairs of methods that produce highly similar distance matrices across 100 subjects’ data. Many of these significant similarities involve the same ROIs with differing thresholding (thr25 vs. WTA), and are commonly found between reconstruction methods (CSD vs DTI). This suggests that the pairwise variation in center-of-gravity coordinates could be ROI specific and less affected by parcellation strategy. These significant similarities are more commonly observed for the CIT168-RL and DISTAL atlases, than for the ATAG atlas. The x, y, and z coordinates for each approach, sorted by ROI, are visualized with boxplots in Figure 5B. These boxplots show how the coordinates are generally within a similar range, with relatively few outliers (with outliers determined by being 1.5 times outside of the interquartile range).

**Figure 5.**
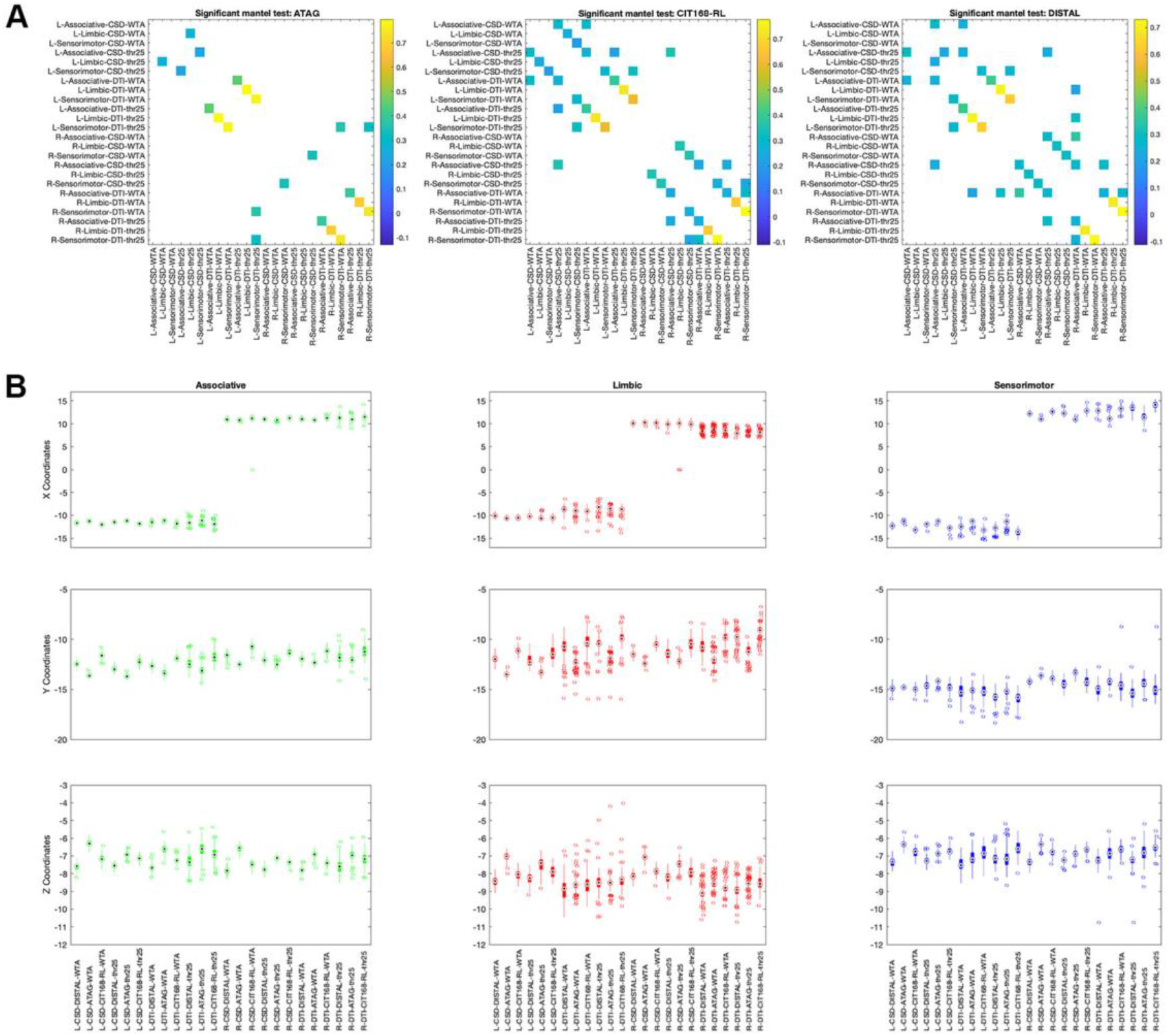
A) The results of Mantel tests to comparing the Euclidean distances between subjects, for each parcellation approach; only significant correlations are shown (α<0.01) B) Boxplots visualizing the x, y, and z coordinates for each approach (central marking: median; bottom and top edges of box: 25^th^ and 75^th^ percentiles; outliers indicated as dots).

Using the TRT dataset and a permutation testing approach, we were able to test the reliability of the center-of-gravity coordinates for each method across scanning sessions. Of the 72 methods tested, 64 displayed significantly lower median distance ranking than as expected by chance (α<0.01). Only 8 parcels were not found to be significantly reliable by this approach; all of them were limbic parcels and obtained with DTI signal modeling.

### 4. Discussion

The present work, by performing STN connectivity-based parcellation, compares four different parcellation methods: one using CSD tractography, and the other using classical single-fiber DTI tractography, both with and without a winner-takes-all parcellation approach. Our results show that all the four different parcellation approaches identified structurally segregated parcels in the human STN.

Our results are in line with the growing body of literature based on non-human primates anatomical and behavioral evidence (Hamani et al., 2017; Karachi et al., 2005, 2009), by showing a well-characterized topographical organization of connectivity in the human STN, with the ventromedial STN prominently connected to limbic cortical areas, dorsomedial STN with associative cortical regions, and dorsolateral STN with sensorimotor cortices. This organization has also been investigated in different human studies using advanced MRI and neuroimaging techniques (Table 2).

**Table 2.** Connectivity-based parcellation of the STN: an overview. Synopsis of the previously existing published works which have attempted the reconstruction of STN functional territories using tractography. For each study the following details are listed: the first author, year of publication, number and type of human subjects, field strength of the MRI scanner used, the number of parcels obtained, the signal modeling and tractography technique used, the method used for STN identification and the clustering protocol.

| *Authors, year* | *Subjects* | *Field Strength* | *Number of parcels* | *Signal modeling* | *Tractography* | *STN identification* | *Clustering method* |
| --- | --- | --- | --- | --- | --- | --- | --- |
| *Lambert et al, 2012* | 12 healthy subjects | 3T | 3 (motor, limbic, associative) | Multi-fiber model (bedpostX) | Probabilistic | Manual delineation on R2* images | Hierarchical clustering |
| *Brunenberg et al, 2012* | 8 healthy subjects | 3T | 3 (Primary motor cortex, supplementary motor area, precentral gyrus) | Q-ball imaging | Probabilistic | Talairach (stereotaxic) | No explicit clustering method |
| *Accolla et al, 2014* | 13 healthy subjects | 3T | 3 (motor, limbic, associative) | Multi-fiber model (bedpostX) | Probabilistic | Morel atlas (histological) | Winner-takes-all |
| *Avecillas-Chasin et al., 2015* | 6 patients (PD) | 3T | 1 (motor) | Single-fiber DTI (StealthViz) | Deterministic | Manual delineation on FLAIR images | No explicit clustering method |
| *Petersen et al., 2017* | 5 patients (PD) | 3T | 1 (motor) | Single fiber DTI; Multi-fiber model (MrTrix) | Deterministic; Probabilistic | Manual delineation on T2 images | No explicit clustering method |
| *Plantinga et al, 2018* | 17 patients (PD) | 7T | 4 (motor, limbic, associative, other) | Multi-fiber model (bedpostX) | Probabilistic | Manual delineation on T2 images | 25% connectivity threshold; no explicit clustering method |
| *Ewert et al., 2018* | 32 healthy subjects | 3T | 3 (motor, limbic, associative) | Diffusion spectrum imaging | Global tractography | Template-based | Winner-takes-all |

Earlier reports identified a posterior-anterior gradient in STN-motor cortical connectivity, where voxels in the posterolateral STN show higher structural and functional connectivity to motor areas and voxels in the antero-medial STN show higher functional connectivity to limbic areas (Brunenberg et al., 2012). Lambert and colleagues used a partially automated clustering method on probabilistic tractography data from 12 healthy subjects and subsequently mapped the connectivity of each parcel to basal ganglia and cortex, revealing a tripartite subdivision of STN that broadly supports findings in animals (Lambert et al., 2012). Similar results were obtained by Accolla and colleagues by mapping cortical connectivity of the STN using an hypothesis-driven approach; the authors also found differences in regional covariance of some brain tissue properties (myelination and iron content) between limbic-associative STN and motor STN, by using quantitative structural MRI (Accolla et al., 2014). Plantinga et al. combined 7T structural MRI and probabilistic tractography to identify the motor domain of STN on 17 idiopathic PD patients before undergoing DBS treatment. The STN was subdivided into four compartments (associative, limbic, sensorimotor and the remaining cortex) and average volumes of each parcels in relation to the whole STN volume are reported (Plantinga et al., 2018). Finally, a tractography-based parcellation based on a group normative connectome of 32 healthy subjects obtained using diffusion spectrum imaging (DSI) paired with a global tractography algorithm (Horn and Blankenburg, 2016) was implemented on a template-based manual delineation of STN and included in a popular standard-space DBS atlas: parcellation was conducted using a WTA approach based on the tripartite model of STN connectivity (associative, limbic and sensorimotor parcels) (Ewert et al., 2018). In agreement with their findings, we described a similar overall topographical parcel organization at the group level, and similar parcel volumes at subject level, with larger volumes for sensorimotor and associative parcels and smaller limbic parcels.

Moreover, we demonstrated that this general topographic organization is preserved regardless of the subcortical atlas used for STN segmentation and of the parcellation pipeline employed; this finding is further corroborated not only by visual inspection but also from the correlation found between center of gravities of parcels obtained from different pipelines, which suggest that the pairwise variation in center-of-gravity coordinates may be only minimally affected by different parcellation strategies. On the other hand, we found that sizes and volumes of each parcel can significantly vary across different methods. In particular, our results suggest that the volume ratio of each parcel is strongly influenced not only by atlas selection (i.e. parcels obtained with a certain atlas tend to have different volumes compared to parcels obtained with another atlas) and pipeline selection (i.e. parcels obtained with a certain pipeline also are systematically different from parcels obtained with another pipeline), but that a highly significant interaction effect may be also found, i.e. parcel volumes obtained with each combination of atlas and parcel tend to behave in a different way. Caution should then be used in interpreting parcel volumes obtained using tractography-derived parcellation as biologically meaningful estimates of the volume of each STN functional territory.

To the best of our knowledge, this is the first work that uses DTI signal modeling combined with probabilistic tractography to reconstruct the entire topographical organization of human STN. Compared to CSD and other complex, multi-fiber modeling techniques, DTI is faster and requires less computing power, making it more suitable for clinical context (Avecillas-Chasin et al., 2015; Berman, 2009; Essayed et al., 2017). A previous study implemented DTI and deterministic tractography-based parcellation in a neuronavigation system to parcellate STN in 6 patients undergoing DBS for PD, but was aimed at identifying only the motor compartment (Avecillas-Chasin et al., 2015). Another pilot study on 5 PD patients compared the reconstruction of motor projections of cortico-STN pathway using CSD and DTI, and found substantial differences in position and shape between these two techniques; they also report higher variability in COG of reconstructed parcels for DTI compared to CSD, suggesting that DTI may provide less reproducible results (Petersen et al., 2017). In line with their findings, parcels obtained with DTI show appreciable differences when compared with those obtained with CSD. The most salient differences were found for the limbic parcel, as these parcels were not reconstructed in a large number of subjects, independently of the atlas employed for STN segmentation. These differences may be due to well-known DTI limitations, in particular when dealing with complex fiber configurations such as fanning and crossing fibers, or regions of high intra-voxel inhomogeneity in fiber orientation (O’Donnell and Westin, 2011) that may lead to underestimation of streamlines. On the other hand, a fair agreement can be observed between sensorimotor and associative parcels, that were reconstructed successfully in almost the totality of subjects even using DTI-based methods.

In our reliability analysis of subthalamic parcellations, we evaluated both between-subjects and within-subject similarity, for those subjects with available TRT data. Between-subjects similarity was assessed using two overlap metrics that slightly differ from each other in their meaning and interpretation: TAO may be interpreted as a global estimate of the reproducibility of the whole pipeline, whereas OBL evaluates the similarity of single parcels obtained with the same procedure. While the choice of different atlases for STN parcellation had a likely negligible effect on TAO values, the impact of signal modeling is evident from our results, where marked differences in TAO values can be observed between CSD-based pipelines and DTI-based pipelines.

However, we suggest that the poor reconstruction of limbic parcels with DTI signal modeling may account for these differences in TAO, as can be deduced from the very low OBL values obtained with DTI-based pipelines for these parcels (see Figure 4A). On the other hand, differences in OBL are lower for associative and sensorimotor parcels, where OBL values obtained with CSD-based pipelines are similar to those obtained with DTI-based pipelines. For the associative parcel, in particular, DTI/WTA pipelines yield systematically higher OBL values when compared to CSD/WTA, regardless of the atlas taken into account. This is, however, the only exception in which DTI-based parcellation seems to outperform a CSD-based method. The general trend suggests that DTI-based parcellation pipelines may provide less reproducible results when compared to CSD-based pipelines, both globally (TAO values) and for each single parcel (OBL values).

It is worthwhile to note, however, that associative and sensorimotor parcels yielded higher OBL values in all the pipelines, including DTI-based pipelines, suggesting that these parcels may be successfully identified with fair reliability also by using simpler signal modeling algorithms such as DTI. This proposition is strengthened by the results of within-subjects similarity, that was evaluated in the TRT dataset by using DSC, a very commonly used similarity measure for neuroimaging. Even if all the parcels obtained very high similarity between test and retest scans (DSC > 0.88) with the only exception of DTI-based limbic parcels (that were probably biased by the high number of reconstruction failures, higher than 50% for most of the atlases and sides), statistically significant differences emerged. Although the effective significance of differences driven by atlas choice or by atlas and pipeline interactions could be questioned, due to their inconsistency between parcel types and the relatively small effect sizes, a significant effect of pipeline selection, with substantially high effect sizes, emerged for all parcels. Interestingly, DTI-based pipelines apparently outperformed CSD-based pipelines in terms of test-retest reliability, showing similar or even higher average DSC values.

This was particularly evident when comparing protocols including a WTA hard segmentation as conclusive step and protocols without WTA hard segmentation. Hard segmentation exclusively assigns each voxel in the target ROI to the parcel displaying higher connectivity values (Behrens et al., 2003). Although widely used in literature (Cacciola et al., 2019a, 2019b; da Silva et al., 2017b;Middlebrooks et al., 2018; Middlebrooks et al., 2018), the use of a WTA approach may introduce a bias, by showing only the most connected voxels, and then imposing a stricter parcellation in respect to the anatomical reality. In particular, it has been suggested that STN may be composed of partially overlapping functional territories rather than well-delineated subdivisions (Alkemade et al., 2015; Lambert et al., 2015; Plantinga et al., 2018): the choice of a WTA-free parcellation protocol would then better fit this anatomical scenario. However, the choice of using no explicit parcellation methods may require the selection of a connectivity threshold to filter out the voxels with lower connectivity profiles (Domin and Lotze, 2019; Johansen-Berg et al., 2005; Tziortzi et al., 2014), introducing an additional source of arbitrariness within the parcellation pipeline. In our parcellation pipeline we used the connectivity threshold of 25%, in line with previous works (Plantinga et al., 2018). Differences in between-subjects similarity indices due to different parcellation approach (WTA, no WTA) are less marked than those deriving from different signal modeling (CSD, DTI); in other words, it seems that the choice of parcellation approach has a smaller influence on between-subjects similarity, compared to the choice of diffusion signal modeling. In contrast with this observation, more marked differences were found in within-subject similarity; in particular, CSD-WTA pipeline resulted in lower average DSC values when compared both to CSD-thr25 or DTI-based pipelines. Our results suggest that, when using higher-level diffusion signal modeling algorithms, WTA parcellation may provide less reproducible results when compared to other parcellation methods. In line with this hypothesis, a recent work demonstrated that, using WTA parcellation, most voxels remain part of the same parcel even after extreme corruption of the underlying DWI dataset (random shuffling of diffusion parameters between voxels). The authors suggested that the use of a connectivity threshold may be a possible solution, by excluding from parcellation results the voxels with lower connectivity values, considering that local noise generally decreases the number of streamlines (Clayden et al., 2019). On the other hand, our results show that, when using simpler signal modeling methods, such as DTI, similarity indices are higher when using WTA segmentation. This apparently counterintuitive result may be interpreted considering that DTI-based pipelines, in general, retrieved lower OBL and TAO values, possibly meaning higher variability of tractographic reconstruction. We hypothesize that the stricter constraints imposed by WTA parcellation may in part mitigate this variability, resulting in higher reproducibility.

Some limitations must be taken into account in the interpretation of our findings. The present work evaluates the reproducibility of different parcellation pipelines by assessing similarity between the resulting connectivity-based parcels across different subjects in a sample; in other words, our outcome measure is not able to assess whether between-subject, similarity is an effect of simple anatomical variability of structures of interest, rather than reproducibility of the parcellation pipeline. However, by using the same sample for the entire analysis, we can reasonably assume as constant between-subjects anatomical variability and interpret differences in similarity as mainly due to the effects of parcellation pipeline.

Another important conceptual issue worth mentioning is that higher reproducibility of a given parcellation pipeline does not necessarily imply higher biological accuracy of the obtained parcels. In the present work we apply an atlas-based method for identification of STN in healthy subjects. When dealing with atlas-based segmentation of the STN, it should be kept in mind that, to date, no gold-standard method is available and that different atlases of the STN provide substantial differences in position and size of this small region of interest (Ewert et al., 2018). To account for differences that may derive from arbitrary atlas selection, we tested our pipelines in three recent, state-of-art subcortical atlases including the STN (Ewert et al., 2018; Keuken et al., 2014; Pauli et al., 2018), and we found that the choice of atlas may have a specific impact not only on parcel volumes, but also on some similarity estimates. In addition, atlases based on healthy young subjects may not take into account differences in STN size introduced by age or underlying pathology (Patriat et al., 2020). The aim of the present study was to test the effects of methodological variables in the processing pipelines on STN parcellation results, rather than propose a gold-standard, biologically meaningful parcellation of this structure; therefore, on small clinical cohorts such as those typically used in STN-DBS studies, it is generally advised to identify STN manually on T2-weighted scans. However, manual identification of STN in larger cohorts is time-consuming and the precise identification of its ventral border could be challenging even on high field strength (7T) T2-weighted scans (Bot et al., 2019). Furthermore, every parcellation pipeline is inevitably affected by all the well-known intrinsic limitations of tractography, such as the inability to determine the precise origin/termination of connections at the synaptic level, difficulty in disentangling intra-voxel fiber heterogeneity and susceptibility to possible misestimates of connectivity due to modeling errors (false positives/negatives) (Cacciola et al., 2019a, 2017; Calamuneri et al., 2018; Chung et al., 2011; Jbabdi and Johansen-Berg, 2011; Parker et al., 2013; Rizzo et al., 2018).

Herein, we only tested hypothesis-driven parcellation protocols, which, in comparison with data-driven parcellation methods, are more subject to selection bias, as the number of target regions is selected *a priori* and is not based on intrinsic properties of the given dataset (Eickhoff et al., 2015). We believe that a hypothesis driven approach fits better to the current situation since the employed connectivity-based subdivision of the STN is grounded on a solid anatomical background in animals (Hamani et al., 2017) and has been replicated in humans (Ewert et al., 2018; Plantinga et al., 2018); moreover, hypothesis-driven parcellation provides more easily interpretable results and would be more suitable in a clinical context, not requiring specific computational expertise to be carried out. Finally, we applied a step-wise track generation approach, based on local parameters; this approach is not as robust to noise, imaging artefacts and partial volume effects as global tractography (Jbabdi et al., 2007; Reisert et al., 2011). Despite this, we have opted for traditional, local step-wise tracking since it is faster and more commonly available than global tractography, and we acknowledge that the global approach may have beneficial effects on the overall reproducibility of the parcellation pipelines.

We believe that our results may have clinical relevance for the development of a robust, reproducible protocol for parcellation of the STN, which may be useful for pre-operative targeting in DBS. Indeed, therapeutic stimulation of STN is often affected by well-known behavioral, cognitive and affective side effects, such as impulse dyscontrol (Frank et al., 2007; Hälbig et al., 2009), irritability and agitation (Merello et al., 2009), psychosis (Kimmel et al., 2013), cognitive and executive functioning deficits (Jahanshahi et al., 2000; Funkiewiez et al., 2004; Rothlind et al., 2015), depression and suicidal ideation (Temel et al., 2006; Voon et al., 2008; Aviles-Olmos et al., 2014). Erroneous targeting of neighboring structures and/or of different STN functional territories is a proposed anatomical substrate for such complications (Greenhouse et al., 2011; Hamani et al., 2017; Temel et al., 2005; Tremblay et al., 2015)In addition, it may be reasonably hypothesized that stimulating the sensorimotor portion of STN may provide better clinical outcomes. A study used probabilistic tractography in PD patients after DBS surgery to retrieve connectivity fingerprints from Volume of Tissue Activated (VTA) models of active electrodes and demonstrated that cortico-STN connectivity with motor and premotor cortical areas was predictive of clinical improvement in tremor, rigidity or bradykinesia (Akram et al., 2017). Moreover, in addition to the well-consolidated framework of sensorimotor STN stimulation in PD, DBS of the anteromedial, limbic/associative STN has been proposed as a useful target for treating refractory OCD (Haynes and Mallet, 2010; Polosan et al., 2019). Noteworthy, an emerging line of evidence suggests that beneficial effects of STN-DBS in OCD may be mediated by structural connectivity to ventrolateral prefrontal cortex and dorsal orbitofrontal cortex (Li et al., 2020; Tyagi et al., 2019). Connectivity-based identification of the associative portion of the STN could then be helpful for optimization of presurgical targeting for OCD. Taken together, these results suggest that an accurate and reproducible individualized targeting of STN sub-regions may represent an important clinical innovation and may lead to better clinical outcomes and fewer undesired side effects after surgery. Our results suggest that, when available, higher-level signal modeling algorithms such as CSD may be recommended since it provides more reproducible results; on the other hand, hard segmentation approaches should be avoided since they may lead to misestimation of volumes of the target regions and lower reproducibility of results. Generally speaking, DTI signal modeling provides less reproducible parcellation, but it may supply however acceptable reproducibility for targeting of the sensorimotor or associative STN territories, in particular when coupled with a WTA parcellation. Finally, it is worth noting that our results are derived from very high-quality MRI acquisitions from the HCP repository (Van Essen et al., 2012) and further evidence is needed to extend these conclusions to MRI acquisitions commonly available in clinical practice.

### Conclusions

The present work tested four different CBP pipelines for reconstruction of STN functional territories. We showed that, regardless of the chosen pipeline, each parcellation provides similar results in terms of location of the identified parcels, but with significant variations in size and shape. Our results also suggest that a more reproducible parcellation may be achieved by using advanced diffusion signal modeling algorithms, such as CSD, and by avoiding hard segmentation as final step of the pipeline (no WTA approach). Further studies are warranted to translate our results into a clinical setting and to demonstrate which technique may lead to more biologically accurate results.

## Acknowledgements

Data were provided by the Human Connectome Project, WU-Minn Consortium (Principal Investigators: David Van Essen and Kamil Ugurbil; 1U54MH091657), funded by the 16 NIH institutes and centers that support the NIH Blueprint for Neuroscience Research; and by the McDonnell Center for Systems Neuroscience at Washington University.

## Author statement

DM: Conceptualization, Investigation, Supervision, Writing – original draft; Writing – review & editing; GAB: Conceptualization, Investigation, Writing – original draft; Writing – review & editing, Visualization; JF: Formal analysis, Writing – review & editing; SB: Conceptualization, Investigation, Writing – original draft; Writing – review & editing, Visualization; AQ: Resources, Data curation, Writing – review & editing; GA: Resources, Data curation, Writing – review & editing; AB: Resources, Data curation; AC: Conceptualization, Investigation, Project administration, Supervision, Writing – original draft; Writing – review & editing.

## Data availability

One dataset of 100 unrelated healthy subjects **and one test–retest dataset, which is a subset of the 1,200 individual MRIs,** were used to support the findings of this study. Data were provided by the Human Connectome Project, WU-Minn Consortium (Principal Investigators: David Van Essen and Kamil Ugurbil; 1U54MH091657) and are publicly available at https://www.humanconnectome.org/study/hcp-young-adult/document/1200-subjects-data-release.

